# Antagonistic histone post-translational modifications improve the fidelity of epigenetic inheritance - a Bayesian perspective

**DOI:** 10.1101/2024.05.07.592892

**Authors:** B. N. Balakrishna Prabhu, Aditya Naman Soni, Sibi Raj B. Pillai, Nithya Ramakrishnan

## Abstract

Histone Post-Translational Modifications (PTMs) are important epigenetic marks regulating gene expression. The specific pattern of histone PTMs present over the gene is critical for turning on/off the expression of the corresponding gene. During DNA replication in mitotic cells, the histone PTMs are dislodged from the mother chromatid, ahead of the replication fork, and distributed uniformly at random among the daughter chromatids. Building on our previous work which modelled the inheritance of a single PTM, the current work considers the effect of an additional antagonistic PTM. We model the joint PTM sequence by an appropriate Markov model and the DNA replication fork as a noisy communication channel. The optimal Bayesian sequence estimator is then employed at each of the daughter chromatids to reconstruct the mother PTM pattern. A high-fidelity reconstruction, aided by the enzyme machinery, is shown to be possible in the presence of epigenetic memory. The structural properties derived for the optimal estimator are then verified through simulations, which show the improvement in fidelity of inheritance in the presence of antagonism. This is further validated through recent experimental data.

## 1 INTRODUCTION

Epigenetic markers regulate gene expression without altering the genetic sequences. They are inherited across cell generations in a reasonably stable manner and influenced by external factors such as diet, environment and stress, leading them to be reversible [1]. Histone Post-Translational Modifications (PTMs) are one of the prominent sets of markers whose patterns across the chromatin play significant roles in gene activation or repression [2]. Certain histone PTMs such as H3K27me3 are known to silence the chromatin region in which they are enriched while a few others like H3K36me3 and H3K4me3 are known to activate the underlying gene for transcription [3, 4].

During DNA replication, the nucleosomes are dislodged and split as a tetramer (H3-H4)_2_ and two H2A H2B dimers from the mother chromatin ahead of the replication fork[5, 6]. This parental histone, along with its PTMs are then transferred to one of the daughter chromatids. The choice of the daughter chromatid is postulated to be take place uniformly at random [7–10]. The random placement of the parental histone tetramers among the daughter chromatids is akin to a discrete signal being corrupted by an IID (independent and identically distributed) noise [11]. Thus, a daughter chromatid has two types of nucleosomes immediately after DNA replication - one set that is directly inherited from the mother chromatin with its PTMs, and another set from a histone pool, without any parental PTMs. In our model, we assume that nucleosomes retain their positional information in the daughter strands with reasonable accuracy as suggested by Reveron-Gomez et al [12], a similar position retention is suggested for the nucleosomes with H3K27me3 in Escobar et al [13]. Certain modifications such as H3K9me3 and H3K27me3 are known to serve as templates for stable inheritance of heterochromatin, along with the associated read-write enzymes [5, 14]. From recent evidence such as cryo-EM reconstructions of human PRC2 bound to functional nucleosomes [15], it can be inferred that the spreading of repressive histone PTMs occurs at the nucleosomal level.

Since the patterns of the histone PTMs across the promoter and gene body play a critical role in the transcriptional regulation [16, 17], it is essential that the distribution of histone PTMs in the parental chromatin is replicated with suffcient fidelity in the daughter chromatids, even if they have only half the information from the parental chromatin. The histone PTMs and several histone chaperones are known to interact with one another so as to aid the effective inheritance of the regulating markers in the daughter chromatids, after DNA replication [5, 18, 19]. The newly generated tetramers are modified based on the sequential pattern of modifications directly inherited from the parental chromatin [6, 20].

Our previous work modeled the DNA replication process by a noisy communication channel, where the inheritance of a specific standalone modification pattern was aided by localized interactions amongst nucleosomes [21]. Since difierent histone PTM marks often interact with one other, we ask the question if the inheritance of a specific histone PTM such as H3K27me3 is aided by the presence of another histone PTM that is antagonistic to it (such as H3K36me3). If yes, how much does the presence of an antagonistic mark improve the fidelity of the original mark? This question is addressed here by employing the principles from Bayesian estimation theory.

## 2 SYSTEM MODEL

Along a region of the chromatin, the presence or absence of a histone PTM is defined as a binary 1 or a 0 respectively. We consider two antagonistic histone PTMs (say *A* and *B*), with PTM *A* being the primary mark. Let 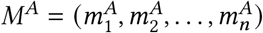 and 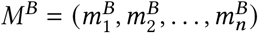 be the binary sequences representing the presence or absence of the histone PTMs *A* and *B* respectively across *n* nucleosomes in the mother chromatin. We _model_ *M*^*A*^ as a first order Markov chain similar to our previous work [21], whereas *M*^*B*^ is modelled as a Hidden Markov Model (HMM) driven by the Markov process *M*^*B*^ .Therefore, the conditional probability for *M*^*A*^ is given by

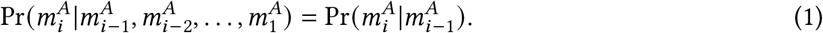

For the HMM *M*^*B*^, the value 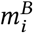 is conditionally independent of others, given 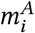.

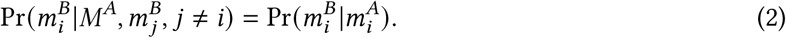

The parameters of the HMM include *α, β* and *μ*, as defined by Fig. 1C. From a biological standpoint, it is better to consider *α* as directly related to the average stretch of 1’s (presence of a mark), while *β* signifies the average stretch of 0’s (absence of a mark).The parameter *μ* represents the probability of the presence of the antagonistic mark (when the primary mark is 0). An example chromatin segment is depicted in Fig. 1A, see Supplementary Information (SI) Fig. S1 for a few other examples. For a single PTM sequence in isolation, the results of our previous work [21] suggest that the inheritance of a histone PTM from the mother to the daughter chromatin has high fidelity when it follows a threshold-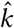 filling algorithm in certain regions of the chromatin (long islands of 1s interspersed by long islands of 0s). In the current work, we analyse how the presence of an antagonistic modification (PTM *B*) in the same region of the chromatin influences the fidelity of the histone PTM *A*’s inheritance.

**Fig. 1.**
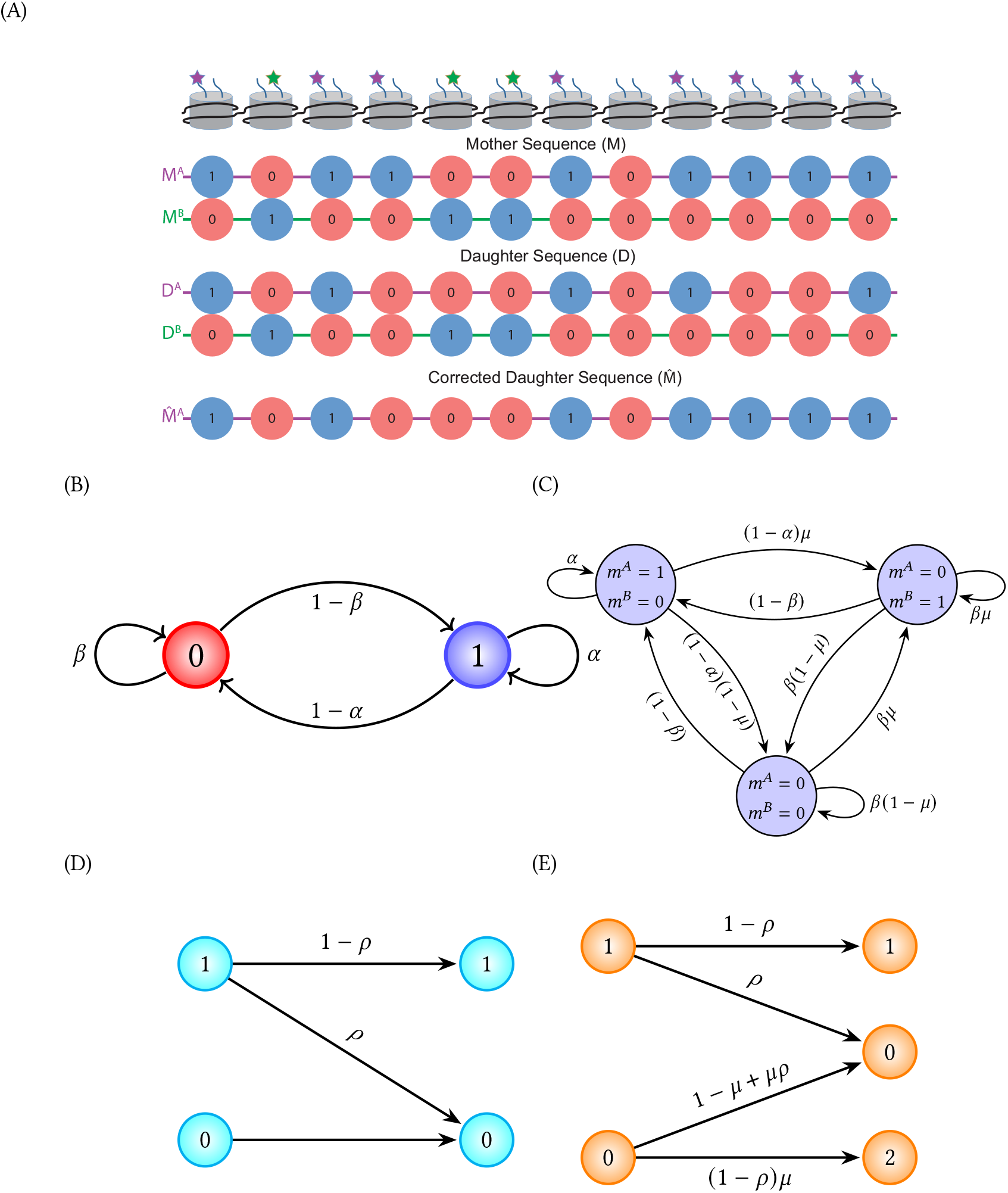
Model Diagrams: (a) The schematic representation of the antagonistic model where 1s and 0s represent the presence and absence of modifications. (b) State diagram for the case considering only PTM-. (c) State diagram for the case considering antagonistic modifications PTM- and PTM-*B* assuming non-bivalent chromatin, i.e. 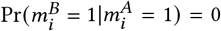 (d) Z-channel model for mother-daughter relationship for a single modification (e) Binary Erasure Channel (BEC) model for mother-daughter relationship with antagonistic modification.

The state transition diagram of the mother chromatin for histone PTM *A* alone is represented by Fig. 1B. This original state diagram is extended to include PTM *B* and represent the transitions in Fig. 1C. Here the states 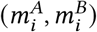 takes values from the set {(0, 0), (0, 1), (1, 0)}. In this manuscript, we make the assumption that the chromatin that is modeled for inheritance is not bivalent [22], i.e., we do not consider antagonistic modifications *A* and *B* to be present in the same nucleosome. Hence (1, 1) is not a permissible state. As can be seen from Fig. 1C, in addition to the transition probabilities *α* (probability of staying in a modified state) and *β* (probability of staying in an unmodified state), a new parameter *μ* is defined as the probability of histone PTM *B* being present, given that histone PTM *A* is absent in the same nucleosome. That is, 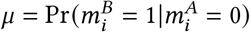.Notice that 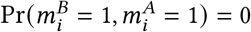 and 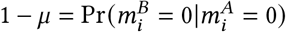.The transition probability 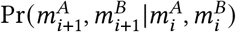 is computed as follows:

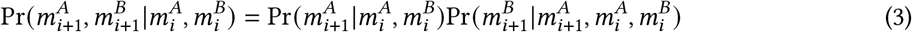

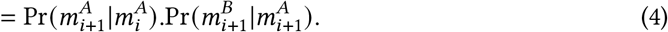

Further details of the values of the transition probabilities are provided in the SI Sec. 1.

During replication, there is a two-fold dilution of the parental histone PTMs in each sister chromatid, which is compensated by the addition of new nucleosomes (without the parental marks), maintaining the nucleosome density [23, 24]. For this study, modifications *A* and *B* are considered to represent the histone PTMs in the same nucleosome, for eg., *A* could be H3K27me3 while *B* could be H3K36me3. While the above pair is a natural choice from the same histone type (H3), one could potentially consider antagonistic pair of PTMs from H3 and H4 in this model. From previous experiments [25, 26], it can be understood that these marks are antagonistic in functionality, i.e., H3K27me3 is a repressive histone PTM as opposed to H3K36me3 which is an activating one [4]. The presence of both of these marks together is rare, but not impossible and can be seen in ‘poised’ regions of the chromatin [27]. It has been suggested by Streubel *et al* [28], that PRC2 and NSD1 (required for H3K27me3 and H3K36me3 deposition respectively) cooperate to mediate their respective marks by a mechanism that is coupled with DNA replication. Our study explores how the Antagonistic histone post-translational modifications improve the fidelity of epigenetic inheritance - a Bayesian perspective additional information from the antagonistic marks in the parental nucleosomes aids in the establishment of the pattern of the primary mark, immediately post replication.

### 2.1 Modeling the replication with antagonistic histone PTMs

In mitotic cells, the split (H3-H4)_2_ tetramers from the mother chromatin are placed in each of the daughter chromatids with equal probability with the help of histone chaperones such as MCM2 [29, 30]. There exist studies showing that both symmetric and asymmetric inheritance (where one daughter chromatid gets a larger share of the parental nucleosomes than its sister) are observed in difierent cell lines [31], even in mESCs [32]. Additionally, symmetric inheritance is significantly hampered in cases where MCM2 is mutated [29]. We model the parameter for distribution between the daughter chromatids as *ρ* and analyze symmetric inheritance (*ρ* = 0.5) from the mother to the daughter chromatids, in this work.

The DNA replication fork is modelled as a noisy communication channel in which one of the two daughter chromatids is given a fraction (1 − *ρ*) of the parental histones, with the remaining newly assembled from a histone pool. For the single modification case, the Z-channel model in Fig. 1D can be used to depict the _DNA replication fork, where the daughter sequence 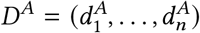_ is obtained as 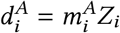,where *Z*_*i*_ ∈ {0,1}is IID with Pr(*Z*_*i*_ = 0) = *ρ*. The process of reconstruction of the mother-like pattern from the daughter is akin to estimating the original signal from the corrupted signal. The additional presence of antagonistic modifications allows us to make a more informative choice during the estimation process. Labelling (0, 0), (1, 0) and (0, 1) by 0, 1 and 2 respectively, the daughter sequence can be represented as 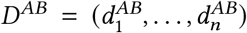 with 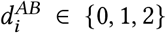. The model with antagonism can then be expressed using 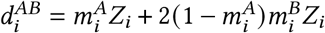 as the daughter observations. The input-output relation is graphically shown in Fig. 1E. The latter model is called the Binary Erasure Channel (BEC) in information theory. Thus, for 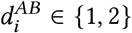, the values of the mother can be correctly guessed from the daughter. In this work we assume that parental nucleosomes preserve their positional order in the daughter chromatids [12]. Though the positioning is subject to some amount of randomness, the effect of this positional jitter is a subject of future work. Let us now estimate the remaining unknown segments of the daughter sequence and demonstrate that the additional antagonistic information is improving the fidelity of inheritance of the primary modification PTM *A*.

### 2.2 Reconstructing the mother chromatin’s pattern from the daughter chromatid

We only need to estimate the mother chromatin’s pattern from the daughter chromatid where 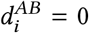 (newly deposited tetramers), between flanking 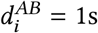 or 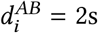 (parental tetramers). We perform a Sequential Maximum A-posteriori Probability (SMAP) based decoding, a Bayesian estimator calculating argmax 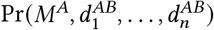 over *M*^*A*^. Notice that

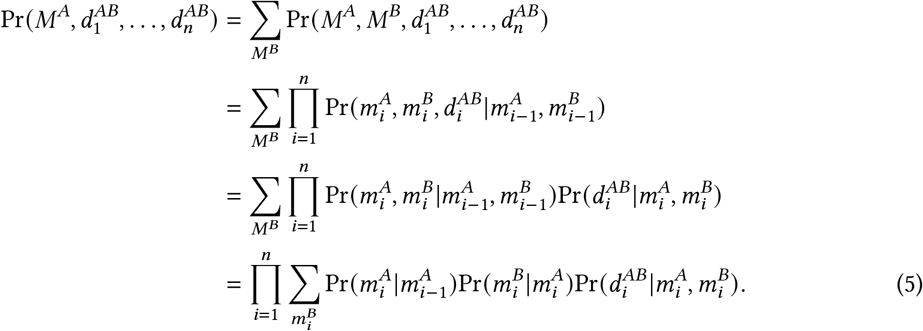

Since the unknown segment is flanked by known values from the mother sequence, for each run of *k:* 0s, there exist four possibilities depending on the flanking values.

Case 1 : {1, 0_:*k*_ 1} - between flanking 1s 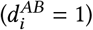 on both sides,

Case 2 : {1, 0_: *k*_ 2} - between a flanking 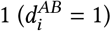 on the left and a 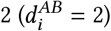 on the right,

Case 3 : {2, 0_: *k*_ 1} - between a flanking 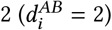 on the left and a 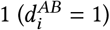 on the right,

Case 4 : {2, 0_: *k*_ 2} - between flanking 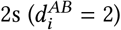 on both sides.

We maximize Eq. (5) using the well known Viterbi decoding algorithm [21] which finds the optimal path that maximizes the joint probability Pr(*M*^*A*^, *D*^*AB*^), from the trellis representing all possible paths. Each of the above four cases will yield a difierent Trellis diagram, this is illustrated in Fig. 2 where the branch metrics are marked on the graph edges. Notice that the branch metrics are evaluated using Eq. (5) for the appropriate states. In each trellis, the winning path is the one with the highest end-to-end path metric.

**Fig. 2.**
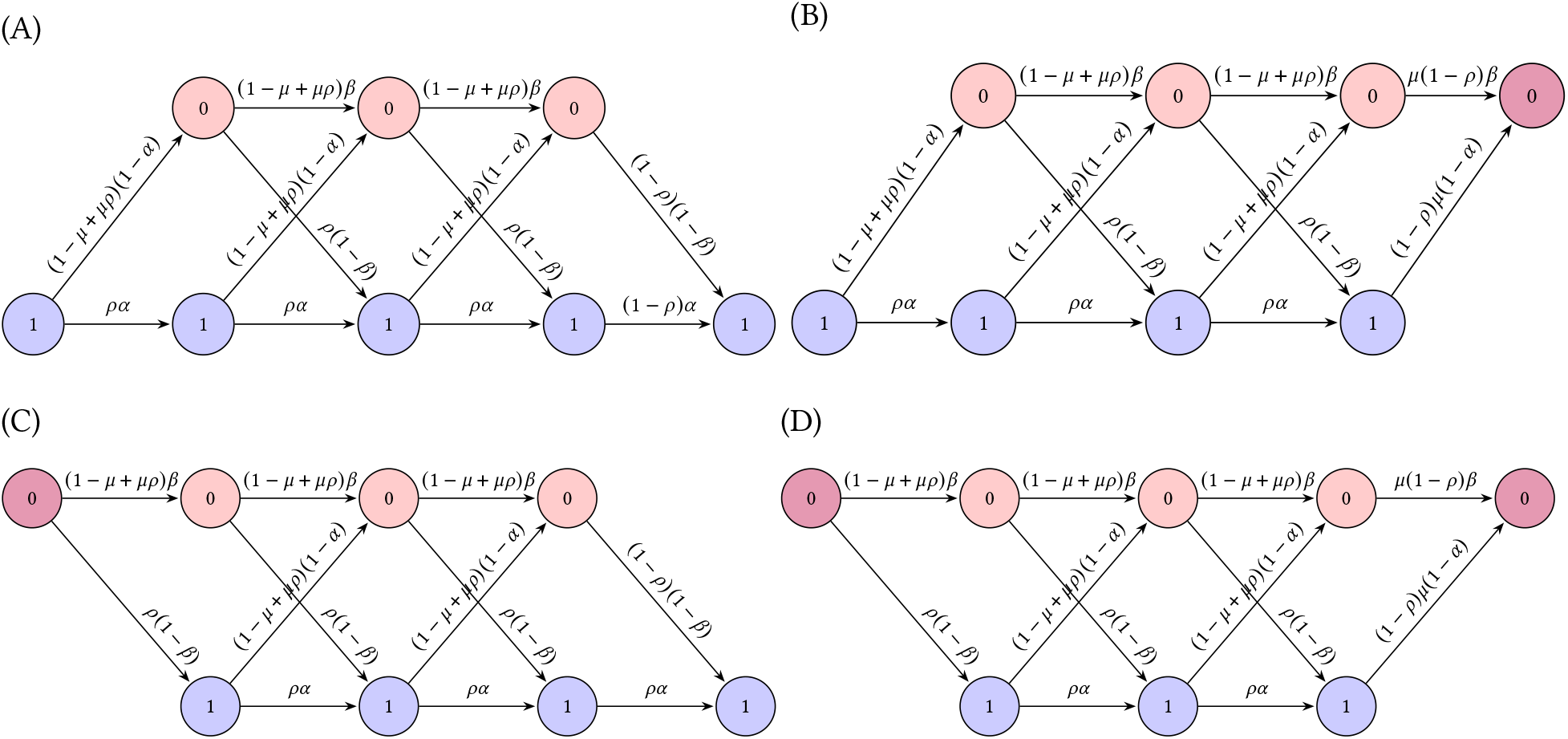
Trellis Decoding: (a-d) Trellis diagrams for the antagonistic modification cases 1-4 respectively as listed in Sec. 2.2 (for 5 nucleosomes). The Viterbi algorithm is used to obtain the reconstructed sequences.

### 2.3 Generalizations of the model

Our model readily generalizes to situations other than the antagonistic case discussed above. Suppose there is a PTM mark *C* instead of the mark *B*, still obeying the Markov relation in (4), with

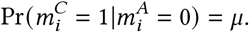

The presence of PTM *C* identifies a parental histone at the daughter strand, and all the marks in that histone should be left unchanged to maintain the fidelity. Therefore, our remaining decision problem is limited to Case 1 – Case 4 listed in Sec. 2.2. Notice that the resulting Trellis diagrams and labels are unchanged during the decoding process, leading to the same estimation performance. As a result, the following models can be easily tackled by our approach.

(1) Non-antagonistic modifications (*M*^*A*^, *M*^*C*^) with 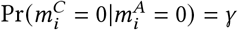 can be accounted by setting μ = 1*− γ* in the trellis.

(2) Nonexclusive antagonistic modifications (*M*^*A*^, *M*^*C*^) with 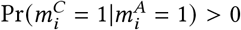 leaves the trellis and performance unchanged. Thus, one can ignore the modification 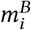 on nucleosomes with both Antagonistic histone post-translational modifications improve the fidelity of epigenetic inheritance - a Bayesian perspective 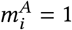 and 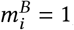,without any loss in performance of estimating the primary modification *“*. Please refer to SI Sec. 1.3 for more details.

(3) Any PTMs in newly deposited histones (such as H3K9me1 [33] and H4K5acK12ac) [34] can be treated as the absence of an antagonistic mark and we proceed with the trellis for decoding.

(4) In cell divisions where the parental histone PTMs are distributed asymmetrically (with a bias towards leading or lagging strands), the parameter *ρ* can account for the aforementioned bias.

From point (2) above, it follows that having a highly correlated non-antagonistic modification can only bring less benefits in reconstructing the primary modification, when compared to the antagonistic case.

## 3 RESULTS

Using simulations of antagonistic sequences, we first obtained the results of the SMAP decoding performed using the Viterbi algorithm. For each *α, β* and *μ* in the interval [0.1, 0.9], binary daughter sequences of length 100 were generated and the corresponding mother sequences were estimated using SMAP decoding. The daughter sequences were obtained from the mother sequences with a *ρ* value of 0.5 i.e., 50% of the mother sequence’s nucleosomes are transferred to one daughter (symmetric inheritance). The results were averaged over 10000 realizations.

_We define the Bit Error Rate (*BER*) corresponding to_ *M*^*A*^ _as the fraction of nucleosomes where the original_ and estimated mother sequences differ with respect to PTM *A*, i.e.,

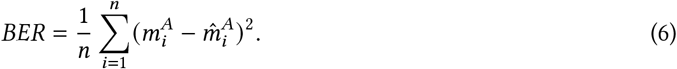

### 3.1 Presence of antagonistic modifications improves the fidelity of PTM ‘s inheritance

We plot the average _*BER*_ as heatmaps for cases of *μ* = 0.2, 0.5 and 0.8 in Fig. 3A, 3B and 3C respectively. The heatmap corresponding to the non-antagonistic (*μ* = 0) case is available in SI Fig. S2a. The Hinton plots in Fig. 3D, 3E and 3F demonstrate the _*BER*_ improvement for *μ* ∈{0.2, 0.5, 0.8} over the non-antagonistic case. From these plots, one can observe that with an increase in *μ*, the average _*BER*_ decreases. This is also evident from the Hinton plots - the number and size of black squares increase with *μ*, indicating that more regions have lower _*BER*_ as compared to the non-antagonistic case.

**Fig. 3.**
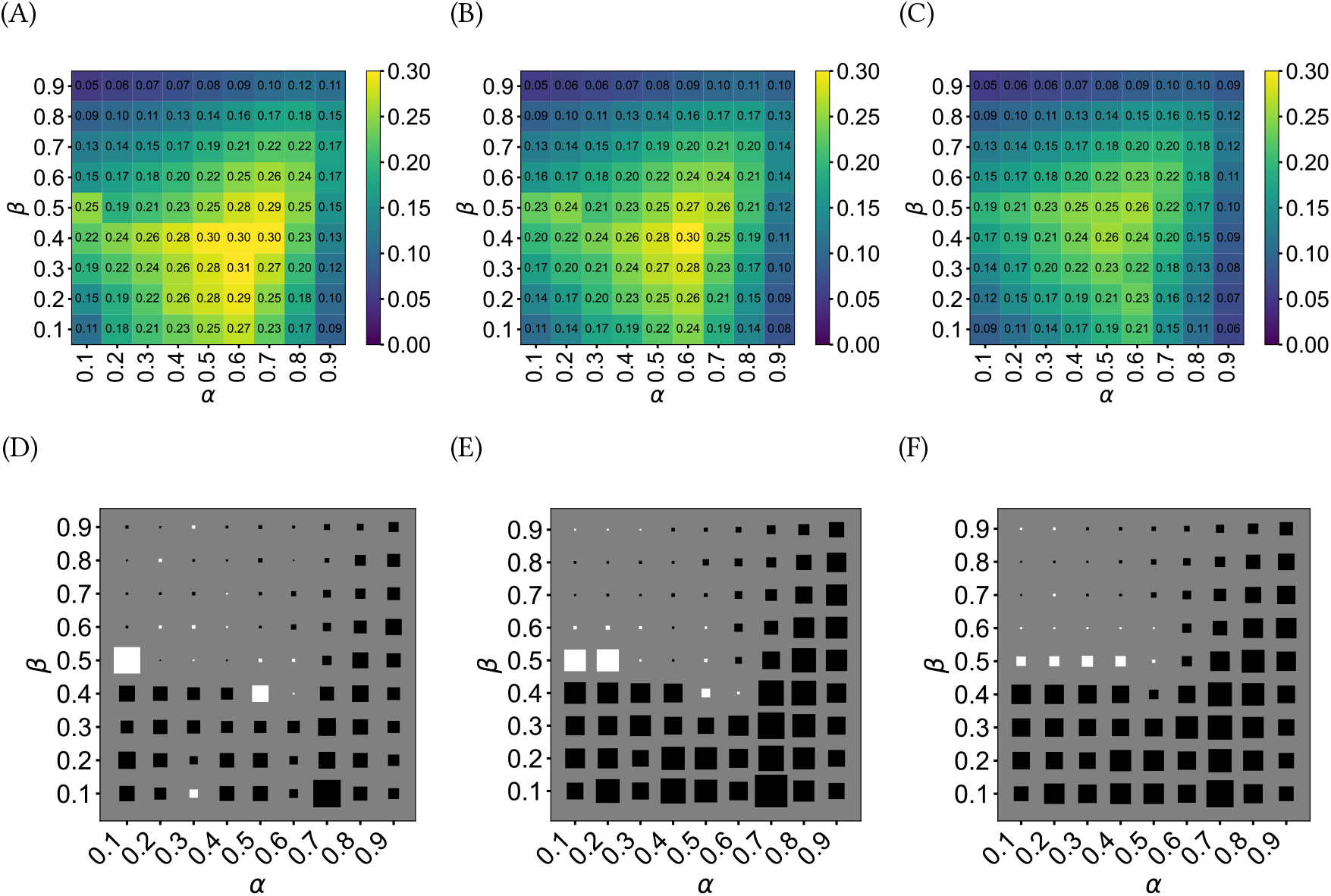
Heatmaps and Hinton plots for simulations: The top row (a, b and c) shows the average _*BER*_ values for the difierent (*α, β, μ*) combinations. The bott om row (d, e and f) shows the corresponding Hinton plots highlighting the difierence between _*BER*_ with no antagonism and difierent levels of antagonism: (a),(d) - *μ* = 0.2; (b),(e) - *μ* = 0.5; (c),(f) - *μ*= 0.8. The heatmap for the non-antagonistic _*BER*_ case is provided in SI in Fig. S2a and is reproduced from our previous work [21]. In the Hinton plots, a black square shows that antagonism lowers the _*BER*_ for that (*α, β*). In the Hinton plots, the largest square has a magnitude of 0.0775 in (d), 0.1152 in (e) and 0.1532 in (f).

Let us discuss the biological relevance of these simulations. We postulate that the recovery for PTM *A* in the daughter chromatid obtained through SMAP decoding could be performed by an ideal enzyme machinery consisting of readers and writers that work in tandem with histone chaperones, transcription factors etc., while catering to the antagonistic modifications. It has to be noted that the estimation of the mother sequence from the daughter sequence requires the knowledge of the mother sequence’s statistical parameters such as *α, β* and *μ*. This is possibly achieved through epigenetic memory effectors such as lncRNA, other PTMs such as H3K4me2, DNA methylation mechanisms and so on [35]. To elucidate how the biological enzymes may behave during cell replication in a realistic scenario, we observe how the Viterbi decoding strategy performs in difierent regions of *α, β* and *μ*. From Fig. 4A, it can be observed that distinct patterns of reconstruction emerge upon applying the Viterbi algorithm. In particular, for Case 1 given in Sec. 2.2, Region A and Region B (marked by blue and white squares respectively in Fig.4A) will obey:

**Fig. 4.**
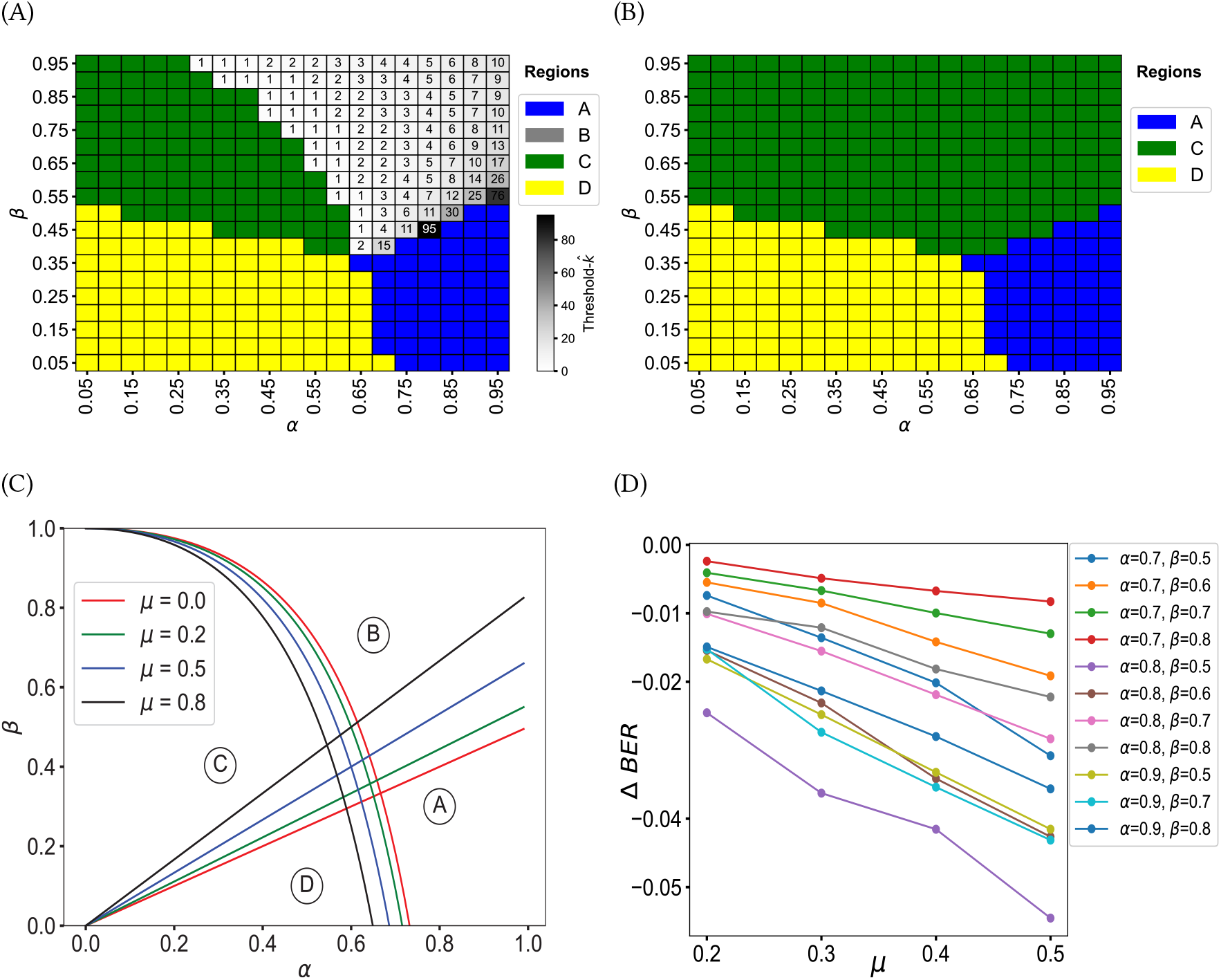
Decoding schemes and BER improvement: Reconstruction patterns for (a) Case 1 for *μ* = 0.2 and (b) Cases 2, 3, 4 for *μ* = 0.2. Threshold-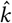: filling is observed only for Case 1 where the value of 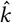: is marked inside the square. The different regions in (a) obey Eq. (S14) and Eq. (S18) in SI Sec. 4. (c) Phase plot indicating the different regions of filling. (d) Δ*BER* between the reconstructed non-antagonistic and antagonistic sequences for different values of *α, β* combinations, across *μ*. The parameters *α, β, μ* are as per Fig. 1C. The simulations were performed for sequences of length 1000, with parameter values *α* ∈ [0.7, 0.9], *β* ∈ [0.5, 0.8] and *μ* ∈ [0.2, 0.6] averaged over 1000 sequences for each combination.

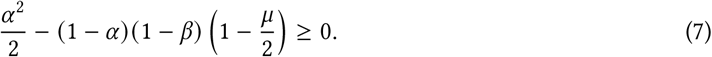

In addition, *α* − (2 − *μ*) *β* ≥ 0 is true in Region A, whereas the reverse inequality is true in Region B. This implies that in Region A, every 0 is converted to a 1 whereas only shorter runs of 0s are modified to 1s in Region B. In Region C (green), the daughter sequence remains unchanged while partial modification is optimal in Region D (yellow).

It has to be noted that except for Case 1, there are only three regions present in the Viterbi reconstruction patterns. Specifically, Region B of Case 1 is replaced by Region C (unfilled 0s) due to the presence of the antagonistic modification in the nucleosome 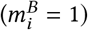 (Fig. 4B).

### 3.2 Higher antagonism leads to more filling of gaps during reconstruction

The threshold-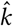: parameter indicating the gap length below which filling will take place in Region B of _Fig. 4A is given by the least integer 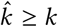,_where

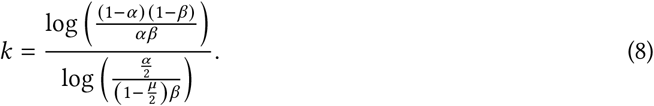

The details of the derivation of threshold-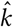 are provided in SI Sec. 4. It can be understood from the above _equation that the threshold-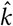_ decreases with *μ*, this can be inferred from Fig. 4C as well. Additionally, from the figure, one could infer that for the chromatin region defined by a high *α* and moderate *β*, a higher level of antagonism may shift the behaviour of reconstruction from Region B (threshold-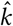 filling) to Region A (zero threshold-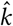) The threshold-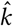 filling is akin to a realistic reconstruction of the mother chromatin’s PTM pattern by the enzyme machinery as the first step in the PTM inheritance process [21].

We then simulated 1000 sequences of length 1000 each, across difierent *α, β* combinations with varying antagonism (*μ*). These sequences were designed to be in Region B of Fig. 4C so that one can analyze the _impact of antagonism on the threshold-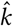_ filling strategy using Eq. (8). The di”erence in BER (*ΔBER*) from the no-antagonism case (*μ* = 0) is shown in Fig. 4D, demonstrating a clear advantage of having the antagonistic modification.

We also simulated 10000 sequences of length 100 each for a positively correlated pair of modifications and observed similar average *BER* to the case where *γ* = 1 − *μ* (SI Fig. S3), as highlighted in Sec. 2.3. The reconstruction patterns for the cases of the trellis in Sec. 2.2 for difierent values of *μ* are provided in SI Fig. S4, S5.

### 3.3 Verification of the model from the experimental data

The ChIP-seq data for the following 5 modifications in Mus *Musculus* was obtained from GSE135029: H3K27me1, H3K27me2, H3K27me3, H3K36me2, H3K36me3. This mESC dataset was chosen since it had both antagonistic modifications (H3K27me3 and H3K36me3) in the same sample, allowing us to verify our antagonistic modifications model. We binarized the data and obtained the sequences for the antagonistic modifications H3K27me3 (repressive modification) and H3K36me3 (activating modification) using the discretization algorithm explained in SI Sec. 3, Fig. S6. An (α, β, μ) tuple was then associated with each nucleosome by considering a window of length 100 around it to calculate the transition probabilities. Further details are provided in SI Sec. 1.

In this data, we focus on chromatin segments with 0.7 ≤ α ≤ 1.0, 0.5 ≤ β ≤ 0.8 and 0.2 ≤ μ ≤ 0.7. A total of 29112 such segments, each of length 100, spread across all chromosomes were identified. These sequences had modifications A (H3K27me3) or *B* (H3K36me3) or none at a specific nucleosome. A small fraction of the total nucleosomes from the data had 1’s in both the modifications A and *B* (5% on average). As pointed out in Sec. 2.3, our results on estimating the primary modification *A* is not affected by ignoring the modification *B* when both are present. We computed threshold-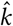 for the daughter sequences (*D*^*AB*^) with and without antagonism (*μ* = 0) using Eq. (8) to facilitate threshold-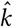 filling so as to yield 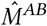.We plot the distributions of the *BER* after averaging over 10 daughter realizations, for the antagonistic and non-antagonistic cases in Fig. 5A and observe that the medians and the interquartile ranges are significantly lower for the antagonistic cases (with p-value 0.001). Fig. 5B shows how the means of the BER distributions vary with *μ*. In SI Fig. S7, the difierence in *BER* for the two cases in a specific region (Chr 18) is shown. These results validate our model suggesting that the presence of antagonistic PTM improves the fidelity of inheritance of the primary PTM.

**Fig. 5.**
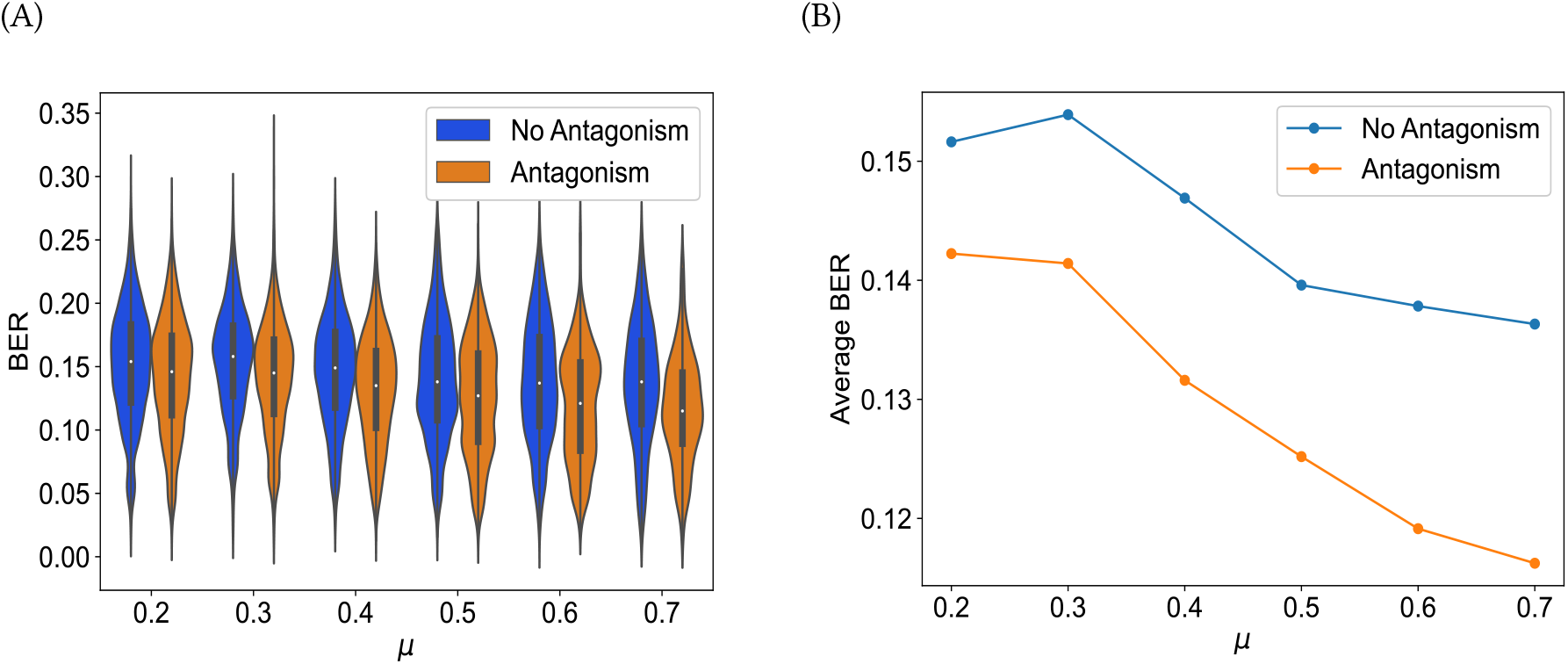
Experimental Validation through the sequences binarized from the ChIP-Seq data from Alabert *et al* [25]: (a) Violin plots for *BER*s obtained for *M*^*A*^ (no antagonism) and *M*^*AB*^ (antagonism) sequences (b) Average *BER’s* for *M*^*A*^ and *M*^*AB*^ sequences. The selected chromatin regions spanned all the chromosomes and had the following statistical parameters: *α* ∈ [0.7, 1.0], *β* ∈ [0.5, 0.8], *μ* ∈ [0.2, 0.7]

## 4 METHODS

The simulations for Fig. 3 were performed on a GPU using Python3 libraries. For each (*α, β, μ*) combination, 10000 mother sequences of length 100 were generated. To obtain *M*^*AB*^_we first generate_ *M*^*A*^ for a given (*α, β*), followed by *M*^*B*^ for a given *μ*. More details are given in SI Sec. 1.1. To correct the daughter sequences that were obtained from the above *M*^*AB*,^ the SMAP decoding algorithm was employed using the Trellis diagrams in Fig. 2 - depending on the boundary conditions. The heatmaps and Hinton plots of Fig. 3 were obtained by comparing the average *BER* (across 10000 sequences) between the original mother and the reconstructed sequence 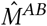.

For verifying our model, the data from GSE135029 (ChIP-seq data from mESC) was discretized to *M*^*AB*^ sequences. The discretization (algorithm explained in SI Sec. 3) was performed on the same GPU using Python3 libraries. The processed bed files from the repository were screened for the presence of H3’s first, followed by the detection of the modifications of interest—H3K27me3 (PTM-*A*) and H3K36me3 (PTM-*B*). We identified 29112 chromatin segments of length 100 with parameters 0.7 ≤ *α* ≤ 1.0, 0.5 ≤ *β* ≤ 0.8, and 0.2 ≤ *μ* ≤ 0.7 (computed from the procedure described in SI Sec. 1.1). The threshold-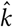 algorithm was performed on this data with threshold-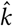 values derived from Eq. (8) for antagonistic (*μ >* 0) and non-antagonistic cases (*μ* = 0). *BER* values were computed for each segment using Eq. (6), the results then averaged over 10 random daughter realizations. The distributions of the computed *BER* values were plotted in Fig. 5A by grouping the sequences over discrete values of *μ*. For each such *μ*, the average *BER* is then computed over all the relevant mother sequences available, this is plotted in Fig. 5B. Statistical significance was tested using a paired Student’s *t*-test (with the usual parameters) by comparing the medians of the two *BER* distributions (antagonistic and non-antagonistic) for every *μ*.

## 5 DISCUSSION AND FUTURE WORK

There have been several models that aim to explain how these epigenetic marks are faithfully transferred from the mother chromatin to the daughter chromatids during DNA replication. Recently, Skjegstad *et al* proposed a 3D polymer configuration model in which they postulate that nucleosomes bind to one another in 3D configurations that affect phenomena such as epigenetic memory and bistability [36]. They employ partial confinement to explain the burst-like nature of certain epigenetic switches. Previously, Dodd *et al* [37] proposed a stochastic model in which cooperation between neighbouring nucleosomes leads to the spreading of modifications over distant regions. Using experimental data from fission yeast, Nickels *et al* [38] suggest that heterochromatin propagates not in a linear manner but as bursts. Alabert *et al* [25] suggest that a domain model (in which methylation states are found in distinct domains) is more likely to explain the antagonism between H3K36me3 and H3K27me3 in mESCs.

Our proposed probabilistic model is not in disagreement with the above theories. While the above models focus on heterochromatin inheritance over long domains, we predict the mechanism (threshold-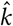 *filling*) by which any modification (particularly silencing modifications) is inherited with high fidelity [21]. In this work, we claim that the antagonistic modifications aid in the inheritance of a given modification as can be seen by the improved fidelity of inheritance. The results from our model show that poised chromatin (with both active and repressive marks in the same chromatin region) are inherited with high fidelity [37]. Prior to our model, Ramachandran and Heinkoff [24] modelled each parental nucleosome assembling at a daughter chromatin as a Bernoulli trial - this is similar to our estimation theory based model. For computing the fidelity, it has to be noted that we consider the case of a single daughter chromatid individually.

The antagonistic roles of H3K36me3 and H3K27me3 have also been widely studied in the past. In Yang *et al* [39], the authors discuss how the antagonistic roles of the aforementioned modifications are necessary for the successful switching of epigenetic states in Arabidopsis for vernalization in winter. In the same work, the authors report a “mirror-like” antagonism between H3K36me3 and H3K27me3 in the nucleation region and gene body. Their model and findings suggest a mutual antagonism between these modifications, similar to those in Alabert *et al*. [25]. Yuan *et al* [40] propose that in HeLa cells, the nucleosomes containing H3K36me3 do not have the H3K27me3 except in newly synthesized H3. The authors also provide evidence that Ash1 (a Trithorax group protein) which is an H3K36-speci!c demethylase, antagonizes PRC2-mediated H3K27 methylation. Some studies point out that marks such as H3K27me3 an H3K36me3 are deposited in the new histones within 2 hours post replication, though not on the same nucleosomes [11, 40]. Our proposed work provides a model for explaining how the presence of H3K36me3 in the neighborhood aids in the faithful copying of the pattern of H3K27me3 from the mother chromatin to the daughter.

There have also been studies on the inheritance of the histone PTMs over several cycles of cell division. Our model not only quantifies the error in inheritance but also postulates the effect of antagonism on the fidelity of inheritance. The molecular mechanisms involved in the the faithful inheritance are not discussed in detail in our work - there could be a plethora of transcription factors, and chromatin remodelers including PRC2 domains, HP1 dimers, RNAi and so on [41].

It is indeed true that writers of histone marks recruit erasers of the marks of opposite functionality (like JMJD2A and JARID for H3K9me2/me3 and H3K4me3 [42]). Even as per our proposed model, the presence of an antagonistic mark causes the primary mark not to be filled during reconstruction of the mother. This will prevent the proximal nucleosomes which lack the modifications (both primary and antagonistic) from getting filled. As shown in our results, this improves the overall fidelity of inheritance of the chromatin mark. Fidelity implies not inheriting when a mark is not present in the mother’s pattern and also inheriting when it is present. The discrepancy in fidelity is measured by the metric *BER* which is the fraction of 0s that have become 1s and 1s that have flipped to 0s from the mother to the daughter. Our model and results suggest that the pattern of repressive marks are maintained better by the presence of antagonistic marks in the neighboring nucleosomes. More broadly, our results suggest that the poised chromatin (where both activating and repressive marks are present in the neighborhood) is inherited with better fidelity, given that the parental marks are symmetrically distributed to the daughter strands. This is in agreement with recent evidence that the regulators of chromatin in poised enhancers are conserved in ESCs [43].

It has been reported that DNA methylation is a critical factor in the faithful inheritance of gene expressions over large timescales [44, 45]. We have not considered the impact of replication-coupled DNA methylation factors in this model at present. We aim to address this concern in our future work.

## DATA AND CODE AVAILABILITY

The primary data analyzed in this study is available at the Gene Expression Omnibus with accession GSE135029. The complete code base is available at the https://github.com/nithya-ramakrishnan-biomodeling/antagonisticviterbi. The corresponding author NR should be contacted for the any other requests.

## ACKNOWLEDGEMENTS

This work was supported by the Department of Electronics, Information Technology, Biotechnology, and Science & Technology of the Government of Karnataka, India. The authors wish to thank Prof.Ranjith Padinhateeri, Dept of Biosciences and Bioengineering, Indian Institute of Technology Bombay for his insightful suggestions.

Antagonistic histone post-translational modifications improve the fidelity of epigenetic inheritance - a Bayesian perspective

## List of Figures

1 Model Diagrams: (a) The schematic representation of the antagonistic model where 1s and 0s represent the presence and absence of modifications. (b) State diagram for the case considering only PTM-*A*. (c) State diagram for the case considering antagonistic modifications PTM-*A* and PTM-*B* assuming non-bivalent chromatin, i.e. 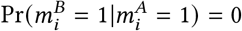. (d) Z-channel model for mother-daughter relationship for a single modification (e) Binary Erasure Channel (BEC) model for mother-daughter relationship with antagonistic modfication. 3

2 Trellis Decoding: (a-d) Trellis diagrams for the antagonistic modification cases 1-4 respectively as listed in Sec. 2.2 (for 5 nucleosomes). The Viterbi algorithm is used to obtain the reconstructed sequences. 7

3 Heatmaps and Hinton plots for simulations: The top row (a, b and c) shows the average *BER* values for the different (*α, β, μ*) combinations. The bottom row (d, e and f) shows the corresponding Hinton plots highlighting the difference between *BER* with no antagonism and different levels of antagonism: (a),(d) - *μ* = 0.2; (b),(e) - *μ* = 0.5; (c),(f) - *μ* = 0.8. The heatmap for the non-antagonistic *BER* case is provided in SI in Fig. S2a and is reproduced from our previous work [21]. In the Hinton plots, a black square shows that antagonism lowers the *BER* for that (*α, β*). In the Hinton plots, the largest square has a magnitude of 0.0775 in (d), 0.1152 in (e) and 0.1532 in (f). 8

4 Decoding schemes and BER improvement: Reconstruction patterns for (a) Case 1 for *μ* = 0.2 and (b) Cases 2, 3, 4 for *μ* = 0.2. Threshold-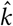 filling is observed only for Case 1 where the value of 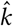 is marked inside the square. The different regions in (a) obey Eq. (S14) and Eq. (S18) in SI Sec. 4. (c) Phase plot indicating the different regions of filling. (d) Δ *BER* between the reconstructed non-antagonistic and antagonistic sequences for different values of *α*, β combinations, across *μ*. The parameters *α,β μ* are as per Fig. 1C. The simulations were performed for sequences of length 1000, with parameter values *α* ∈ 2 [0.7, 0.9], *β* ∈ 2 [0.5, 0.8] and *μ* ∈ 2 [0.2, 0.6] averaged over 1000 sequences for each combination. 13

5 Experimental Validation through the sequences binarized from the ChIP-Seq data from Alabert *et al* [25]: (a) Violin plots for *BER*s obtained for *M*^*A*^ (no antagonism) and *M*^*AB*^ (antagonism) sequences (b) Average *BER’*s for *M*^*A*^ and *M*^*AB*^ sequences. The selected chromatin regions spanned all the chromosomes and had the following statistical parameters: *α* ∈ 2 [0.7, 1.0], *β* ∈ 2 [0.5, 0.8], *μ* ∈ 2 [0.2, 0.7] 14

## Notes

### Competing Interest Statement

The authors have declared no competing interest.

### Summary of Updates

Generalization of the model was performed and updated in Sec 2.3 Aditya Naman Soni was added as the second author

