## Supplementary material for "Antagonistic histone post-translational modifications improve the fidelity of epigenetic inheritance - a Bayesian perspective": SI

### 1 MODEL AND METHODS

In our study, we analyze two binary sequences: $M^A = (m_1^A, m_2^A, \ldots, m_n^A)$ and $M^B = (m_1^B, m_2^B, \ldots, m_n^B)$. Sequence $M^A$ represents a specific histone Post Translational Modification (PTM), while sequence $M^B$ denotes an antagonistic PTM to $M^A$ (Fig. 1A). Our goal is to estimate the transition probabilities $\alpha$ and $\beta$ within sequence $M^A$, and the conditional probability $\mu$ between sequences $M^A$ and $M^B$. Referring to Fig. 1B of the main manuscript, $\alpha$ is the probability that a site remains modified (represented as 1) given that it was modified on the preceding nucleosome. Mathematically, this can be expressed as:

$$\alpha = Pr(m_i^A = 1 | m_{i-1}^A = 1). \tag{S1}$$

Similarly, $\beta$ represents the conditional probability that a site remains unmodified (represented as 0) *i.e.*

$$\beta = Pr(m_i^A = 0 | m_{i-1}^A = 0). \tag{S2}$$

In Fig. 1C we take the parameter $\mu$ as

$$\mu = Pr(m_i^B = 1 | m_i^A = 0), \tag{S3}$$

identically for all $1 \leq i \leq n$. Thus, $\mu$ quantifies the statistical relationship between sequences $M^A$ and $M^B$, specifically the likelihood of observing an antagonistic modification (in $M^B$) given the absence of modification (in $M^A$).

For the case of nonexclusive antagonistic modifications $A$ and $C$, we define $\gamma$ as follows:

$$\gamma = Pr(m_i^B = 0 | m_i^A = 0) = \gamma = Pr(m_i^B = 1 | m_i^A = 1). \tag{S4}$$

#### 1.1 Computation of $\alpha$, $\beta$, $\mu$ and $\gamma$

Given a pair of binary sequences $M^A = \{m_1^A, m_2^A, \ldots, m_n^A\}$, and $M^B = \{m_1^B, m_2^B, \ldots, m_n^B\}$, we define the transition counts necessary for estimating $\alpha$ and $\beta$:

$$n_{11} = \text{Number of transitions from } m_i^A = 1 \text{ to } m_{i+1}^A = 1,$$
$$n_{10} = \text{Number of transitions from } m_i^A = 1 \text{ to } m_{i+1}^A = 0,$$
$$n_{01} = \text{Number of transitions from } m_i^A = 0 \text{ to } m_{i+1}^A = 1,$$
$$n_{00} = \text{Number of transitions from } m_i^A = 0 \text{ to } m_{i+1}^A = 0.$$

Authors' address: B. N. Balakrishna Prabhu; Aditya Naman Soni; Sibi Raj B. Pillai; Nithya Ramakrishnan.

The probabilities $\alpha$ and $\beta$ for sequence $M^A$ are then calculated as follows:

$$\alpha = \frac{n_{11}}{n_{11} + n_{10}}, \tag{S5}$$

$$\beta = \frac{n_{00}}{n_{00} + n_{01}}. \tag{S6}$$

**Estimating $\mu$ and $\gamma$ from $M^A$ and $M^B$**

The probability $\mu$ is given by:

$$\mu = \frac{\sum_{i=1}^{n} \mathbb{1}_{\left(m_i^A=0 \text{ AND } m_i^B=1\right)}}{\sum_{i=1}^{n} \mathbb{1}_{\left(m_i^A=0\right)}}, \tag{S7}$$

where $n$ is the length of the sequences, and $\mathbb{1}_{(\cdot)}$ is the indicator function, yielding 1 if the condition inside is true, and 0 otherwise. For the case of positively correlated pair of PTM's the probability $\gamma$ is given by:

$$\gamma = \frac{\sum_{i=1}^{n} \mathbb{1}_{\left(m_i^A==m_i^B\right)}}{\sum_{i=1}^{n} \mathbb{1}_{\left(m_i^A\right)}}. \tag{S8}$$

### 1.2 Derivation of steady-state probabilities

We represent the joint sequence $M^{AB} := \{(m_i^A, m_i^B), 1 \leq i \leq n\}$ as a Markov model and derive the steady-state probabilities. From the state diagram (Fig. 1C), we consider the transition probability matrix representing the 3 states $S$, $T$ and $U$ as follows:

| | $S$ | $T$ | $U$ |
|---|---|---|---|
| $S$ | $\alpha$ | $(1-\alpha)\mu$ | $(1-\alpha)(1-\mu)$ |
| $T$ | $(1-\beta)$ | $\beta\mu$ | $\beta(1-\mu)$ |
| $U$ | $(1-\beta)$ | $\beta\mu$ | $\beta(1-\mu)$ |

where $S = (m_i^A = 1, m_i^B = 0)$, $T = (m_i^A = 0, m_i^B = 1)$ and $U = (m_i^A = 0, m_i^B = 0)$. The steady-state probability vector, denoted by $\left[\Pi_S, \Pi_T, \Pi_U\right]$, obeys

$$\begin{bmatrix}\Pi_S & \Pi_T & \Pi_U\end{bmatrix} \cdot \begin{bmatrix} \alpha & (1-\alpha)\mu & (1-\alpha)(1-\mu) \\ (1-\beta) & \beta\mu & \beta(1-\mu) \\ (1-\beta) & \beta\mu & \beta(1-\mu) \end{bmatrix} = \begin{bmatrix}\Pi_S & \Pi_T & \Pi_U\end{bmatrix}. \tag{S9}$$

Using $\Pi_S + \Pi_T + \Pi_U = 1$ and solving the balance equations, we get the steady-state probabilities as follows:

$$\Pi_S = \frac{1-\beta}{2-\alpha-\beta}, \tag{S10}$$

$$\Pi_T = \frac{(1-\beta)(1-\alpha-\beta)\mu}{2-\alpha-\beta} + \beta\mu, \tag{S11}$$

$$\Pi_U = \frac{(1-\beta)(1-\alpha-\beta)(1-\mu)}{2-\alpha-\beta} + \beta(1-\mu). \tag{S12}$$

Here, $\Pi_S$, $\Pi_T$ and $\Pi_U$ represent the stationary probabilities corresponding to the presence of PTMs $A$ and $B$ in a nucleosome.

#### 1.3 Non-exclusive antagonistic modifications

In the mathematical model for the system, we considered two antagonistic pairs which are mutually exclusive, i.e., the possibility of both the antagonistic marks ($A$ and $B$) being simultaneously present in a nucleosome was neglected. Let us now describe how this model and the results can be generalized to the non-exclusive antagonistic model with $\Pr(m_i^B = 1 | m_i^A = 1) = \nu > 0$. Assume that the sequence $(M^A, M^B)$ evolves according to Equation (4) in the main manuscript. After replication, each daughter strand directly acquires the marks from half the parental nucleosomes. As before, our objective is to recover the primary PTM mark $A$ at one of the daughter chromatids. The possible observations at each nucleosome is $(d_i^A, d_i^B) \in \{(0,0), (1,0), (0,1), (1,1)\}$. By calling the state $(1,1)$ as $1'$, we can take $d^{AB} \in \{0, 1, 2, 1'\}$ as the output observation for each daughter nucleosome, see Section 2.2 of the main manuscript.

Notice that if any of the parental mark ($A$ or $B$) is found to be present in the daughter, then no immediate change is necessary at that nucleosome, since the mother PTMs are already revealed. So the estimation and decision rules are only necessary when $d_i^{AB} = 0$. From the PTM $A$ point of view, the decision scheme has to again consider only the four cases listed in Section 2.2 of the main manuscript. The only difference here is that both $d_i^{AB} = 1$ and $d_i^{AB} = 1'$ will lead to the state with $m_i^A = 1$ in the trellis diagrams shown in Fig. 2. However, it can be readily verified that the branch metrics of the trellis remain the same, irrespective of the known state is due to 1 or $1'$. In other words, once $m_i^A$ is revealed to be 1, then the value of $m_i^B$ has no role, and can be ignored for estimating $M^A$. So the model, decoding scheme and the results are independent of the value $\nu$, thus easily accommodating the case of non-exclusive antagonistic models, as can be possibly observed in bivalent chromatin.

### 2 SIMULATION RESULTS

Fig. S1 shows the plots showing the effect of varying parameters $\alpha$, $\beta$, and $\mu$ on the simulated mother sequences. Using the SMAP decoding algorithm we obtain the difference in *BER* between the original mother sequences and the reconstructed mother sequences for $\alpha, \beta \in (0, 1)$ and $\mu = 0$ (non-antagonistic case). The heatmap and violin plot in (Fig. S2) was obtained by averaging over 10000 simulated sequences for each $(\alpha, \beta)$ pair and reproduces the result from our earlier work[1].

Fig. S3 shows the heatmaps of the average *BER* for simulations comparing $\mu$ and $\gamma$ for different values of $(\alpha, \beta)$.

Fig. S4 shows the variation for threshold-$\hat{k}$ values for different $(\alpha, \beta, \mu)$ combinations for Case 1(Sec. 2.2 of the main manuscript).

Fig. S5 shows the variations of the boundaries with $\mu$ for Cases 2, 3, and 4(Sec. 2.2 of the main manuscript.

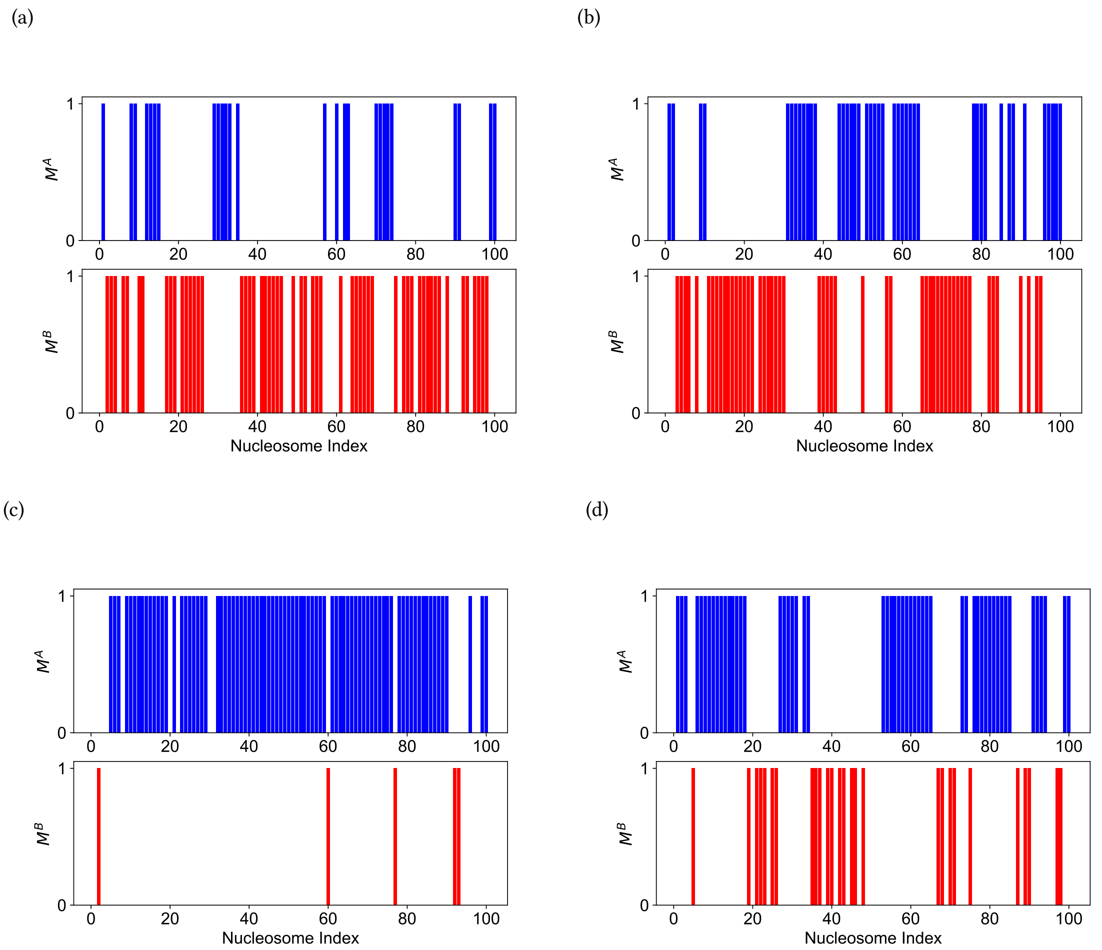


Fig. S1. Simulation Plots: Plots showing the effect of varying parameters $\alpha$, $\beta$, and $\mu$ on the simulated sequences. (a) $\alpha = 0.6$, $\beta = 0.8$, $\mu = 0.8$, (b) $\alpha = 0.8$, $\beta = 0.8$, $\mu = 0.8$, (c) $\alpha = 0.9$, $\beta = 0.4$, $\mu = 0.2$, and (d) $\alpha = 0.9$, $\beta = 0.9$, $\mu = 0.5$.

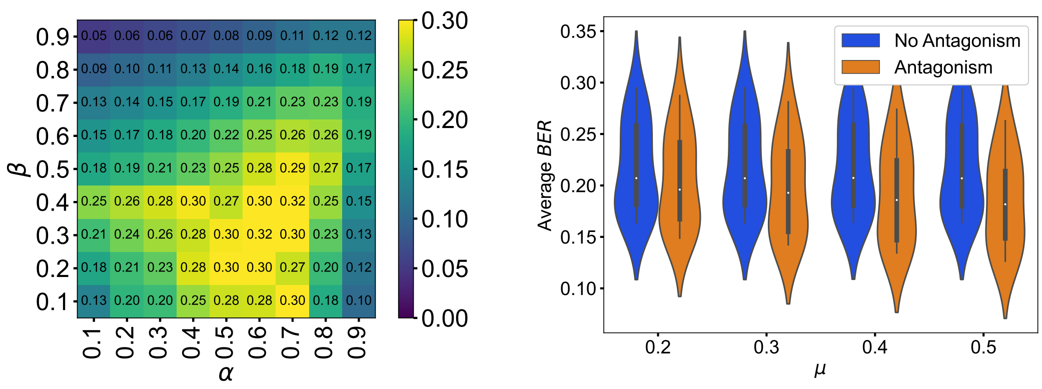


Fig. S2. *BER* in simulations: (a) Heatmaps for the *BER* between the original mother sequence ($M_i^A$) and the reconstructed mother sequence ($\hat{M}_i^A$) obtained using SMAP decoding for sequences without antagonism ($\mu = 0$). (b) Average *BER* of reconstructed non-antagonistic sequences and the antagonistic sequences obtained through simulations averaged over 1000 sequences of length 1000 each, with parameters $\alpha \in [0.7, 0.9]$, $\beta \in [0.5, 0.8]$ and $\mu \in [0.2, 0.6]$.

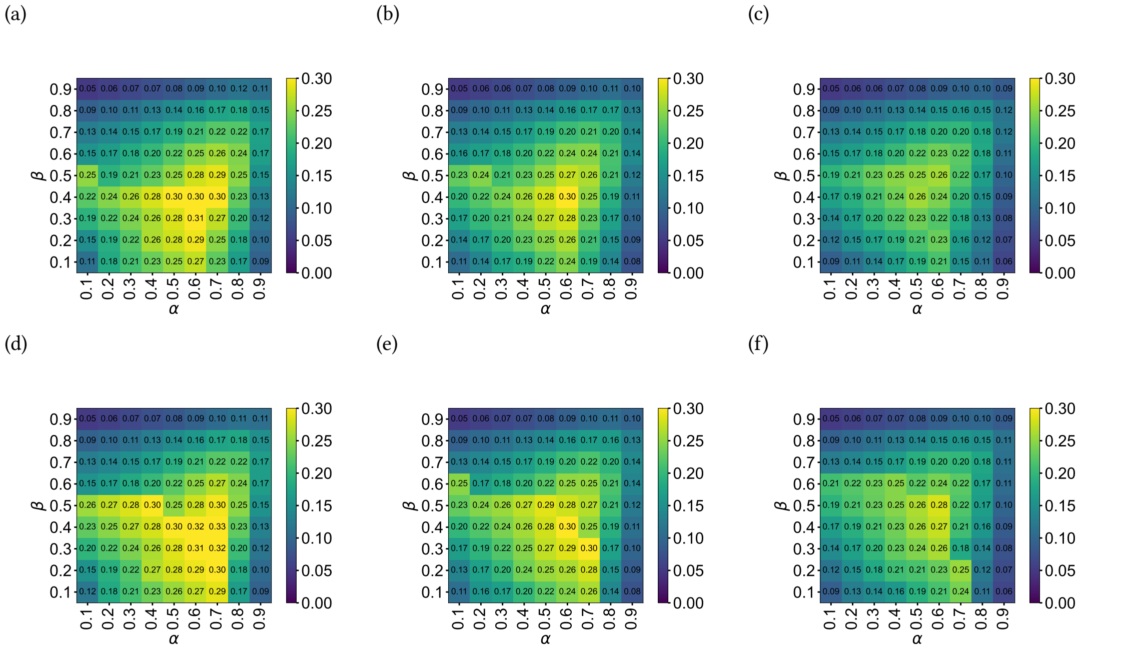


Fig. S3. Generalization to non-antagonistic pair of modifications: The top row (a, b and c) shows the heatmaps for average *BER* values (over 10000 sequences )for the different $(\alpha, \beta, \mu)$ combinations, for the reconstruction performed with the SMAP algorithm. The bottom row (d, e, and f) shows the corresponding plots for $\alpha, \beta, \gamma$ values (non-antagonistic cases). In (a, d - $\mu = 0.2, \gamma = 0.8$, in b, e - $\mu, \gamma = 0.5$, c, f - $\mu = 0.8, \gamma = 0.2$).

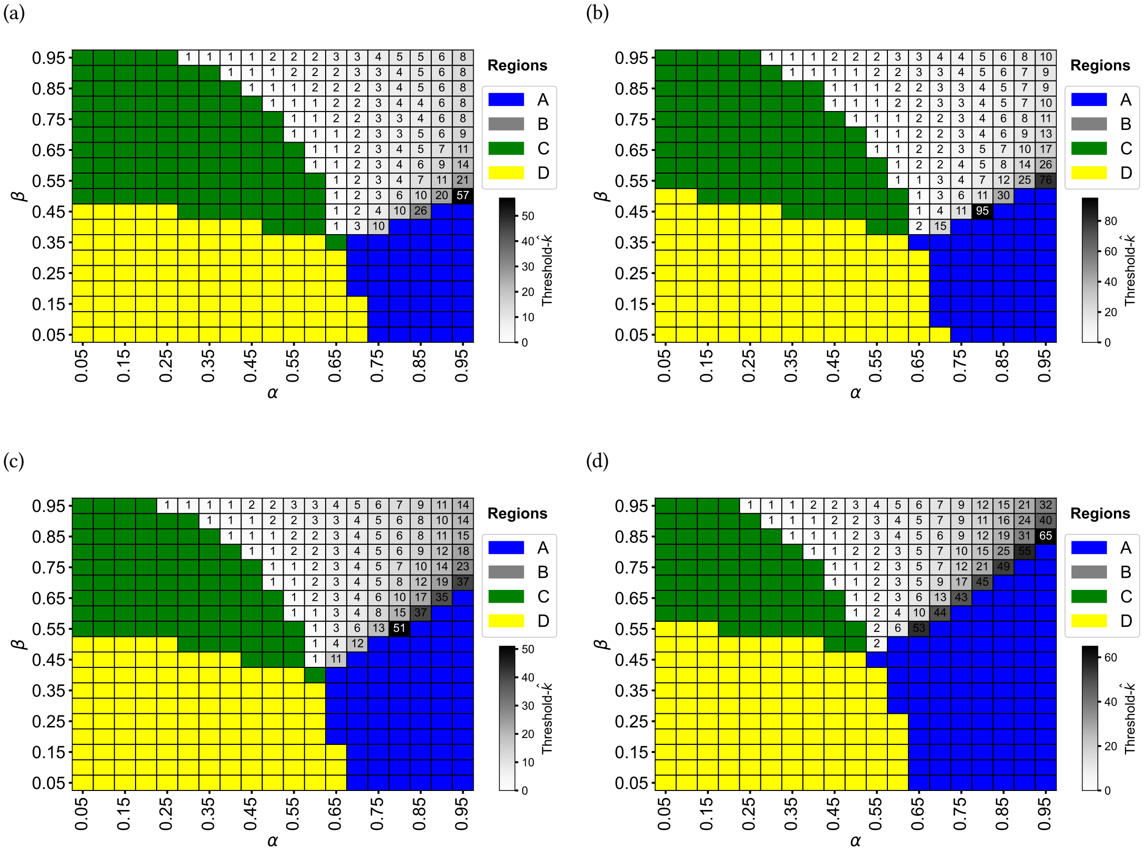


Fig. S4. Threshold-$\hat{k}$ values vary with $\mu$: Reconstruction patterns for (a)$\mu = 0.0$, (b) $\mu = 0.2$, (c) $\mu = 0.5$, (d) $\mu = 0.8$.

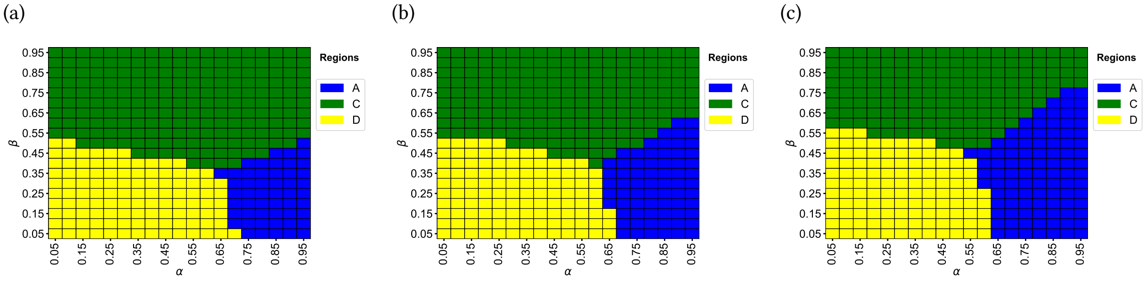


Fig. S5. Region boundaries of reconstruction vary with $\mu$: Reconstruction patterns for Cases 2, 3, 4 (A) $\mu = 0.2$, (B) $\mu = 0.5$, (C) $\mu = 0.8$.

### 3 DISCRETIZATION ALGORITHM

The PTM data from the experiments obtained from [2] included processed bed (browser extensible data) files from the ChIP-Seq experiments. They are transformed into binary sequences that indicate the presence or absence of the PTMs in a nucleosome. The algorithm is detailed in the flowchart of Fig. S6. Binarised versions of PTMs H3K27me3 and H3K36me3 represent sequences $M^A$ and $M^B$ respectively in our work. Additional modifications—H3K27me1, H3K27me2, and H3K36me2 are also considered in detecting the presence of H3.

The PTM data [2] initially exhibits a 500-base pair overlap between consecutive entries, leading to the convolution of the measured enrichment values across the base-pair segments. This is corrected by a deconvolution step, which involves taking the difference over the two overlapping base-pair segments. We start deconvolution from a segment where the enrichment is negligible.

In the base pair segments with negligible enrichment values, to distinguish between the cases where (i) the nucleosome is itself absent or (ii) the PTM is absent in the nucleosome, we use the following approach: If the enrichment values in any of the 5 modifications exceed 0.5 or the sum of enrichments for the 5 PTMs exceeds 1.25, then we consider it the latter case.

Following deconvolution and nucleosome detection, to address the abrupt changes in the enrichment values, we apply a median filter with a window size of 5 nucleosomes for smoothening. The smoothed data is then binarized with a threshold of 0.5.

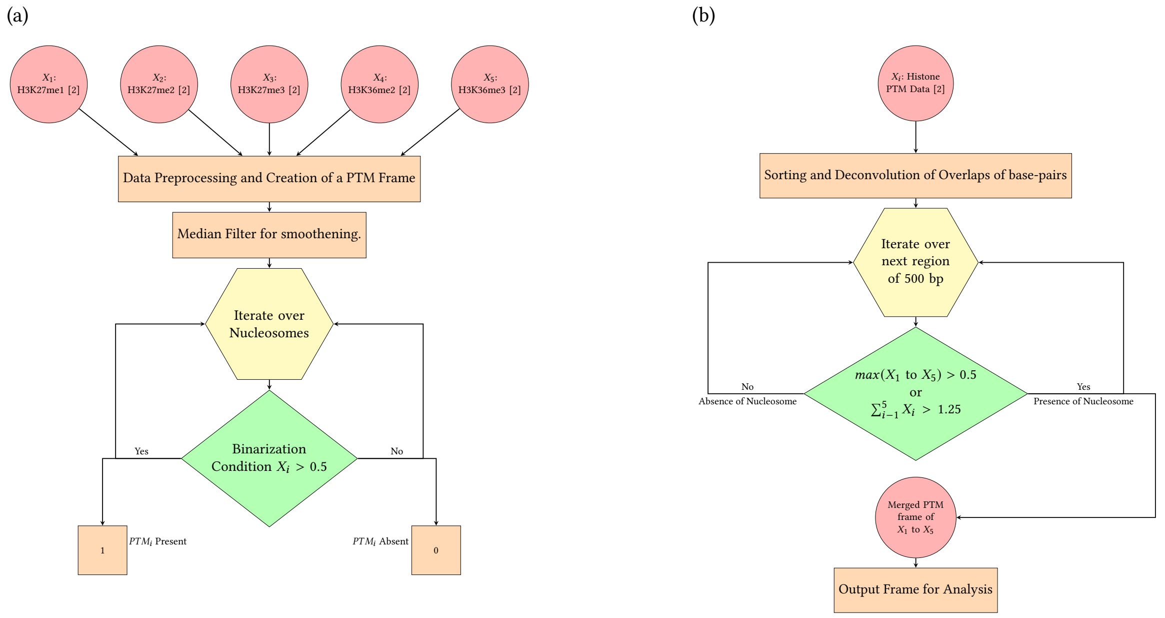


Fig. S6. (A) Discretization algorithm for the binarization of the ChIP-Seq data obtained from the [2]. (B) The pipeline for preprocessing the data includes deconvolution and nucleosome detection.

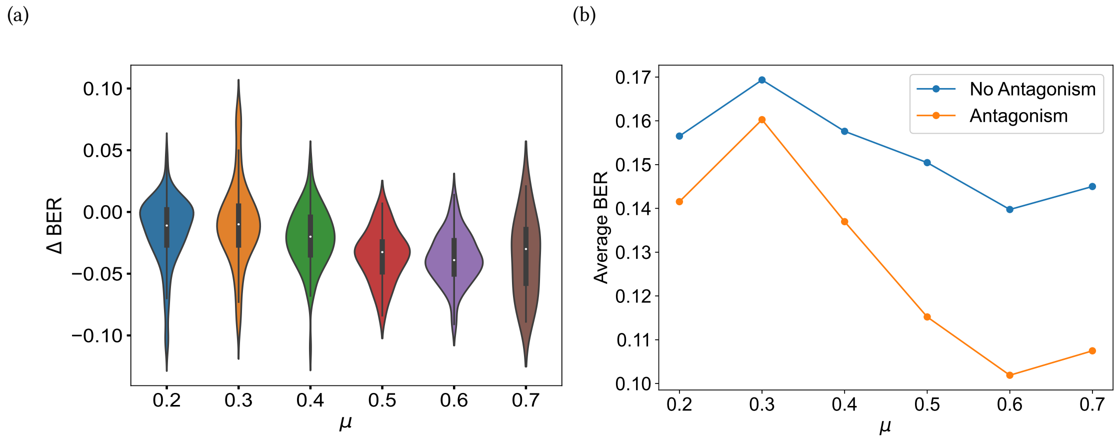


Fig. S7. Experimental Data: (Chr-18) a) Violin plot of the difference in *BER* between the antagonistic and non-antagonistic cases and (b) line plot of the means of *BER* distributions for the non-antagonistic and antagonistic cases for the sequences binarized from the ChIP-Seq data in Chr 18 of the mESCs obtained from [2]. The selected chromatin regions were found to have the statistical parameters: $\alpha \in [0.7, 1.0]$, $\beta \in [0.5, 0.8]$, $\mu \in [0.2, 0.7]$.

#### 3.1 Delta BER Plots

Following the discretization procedure, we plotted the $\Delta BER$ (Fig. S7a) and the means of $BER$ (Fig. S7b) for the antagonistic and non-antagonistic cases for the data in a specific region - Chr-18 of the mESC dataset from [2].

### 4 DERIVATION OF PHASE TRANSITION EQUATIONS

Refer to the trellis diagram in the main manuscript (Fig. 2A). In Case-1, given a gap of $k$ zeros$(1, 0_k, 1)$, if an all-one path has a higher path metric than an all-zero path, we get,

$$\left(1-\mu+\frac{\mu}{2}\right)\cdot(1-\alpha)\cdot\left(\left(1-\mu+\frac{\mu}{2}\right)\cdot\beta\right)^{k-1}\cdot\frac{1-\beta}{2}\leq\left(\frac{\alpha}{2}\right)^{k+1}, \tag{S13}$$

Simplifying,

$$\frac{(1-\alpha)(1-\beta)}{2}\leq\frac{\left(\frac{\alpha}{2}\right)^{k+1}}{(1-\frac{\mu}{2})^{k}\cdot\beta^{k-1}}. \tag{S14}$$

The boundary of the region where the gaps are left unfilled can be evaluated by substituting $k = 1$ in Eq. (S14), to obtain the condition

$$\frac{(1-\alpha)(1-\beta)}{2}=\frac{\left(\frac{\alpha}{2}\right)^{2}}{(1-\frac{\mu}{2})}. \tag{S15}$$

Simplifying, we get

$$\frac{\alpha^2}{2}-(1-\alpha)(1-\beta)\left(1-\frac{\mu}{2}\right)=0. \tag{S16}$$

#### 4.1 Derivation of threshold-$\hat{k}$

Taking logarithm on both sides of Eq. (S14) and simplifying, we get

$$\log\left(\frac{(1-\alpha)(1-\beta)}{2}\right) = k\log\left(\frac{\frac{\alpha}{2}}{\left(1-\frac{\mu}{2}\right)\beta}\right) + \log\left(\frac{\alpha\beta}{2}\right). \tag{S17}$$

Therefore,

$$k = \frac{\log\left(\frac{(1-\alpha)(1-\beta)}{\alpha\beta}\right)}{\log\left(\frac{\frac{\alpha}{2}}{(1-\frac{\mu}{2})\beta}\right)}. \tag{S18}$$

The straight line shown in Fig. 4C corresponds to the denominator in Eq. (S18) being 0. This yields

$$\alpha = (2-\mu)\beta. \tag{S19}$$

### 5 SIMULATIONS OF DIFFERENT CASES OF DAUGHTER CHROMATIN

For the simulations described in Sec. 3 of the main manuscript, we analyse the results for no-antagonism ($\mu = 0.0$) and different ranges of antagonism ($\mu = 0.2, 0.5, 0.8$). The results are shown in Fig. S4 (Case 1) and Fig. S5 (Cases 2, 3, 4). Since Cases 2, 3, and 4 are having similar results, we show a representative case (Case-3) in Fig. S5.

From Fig. S4 one can infer that region A expands with $\mu$. additionally, it can be observed that the threshold-$\hat{k}$ increases with $\mu$ for high values of $(\alpha, \beta)$. As in Fig. 4 of the main manuscript, region B (threshold-$\hat{k}$) can be observed only in Case-1.
